# Time of Sample Collection Critical for Microbiome Replicability

**DOI:** 10.1101/2022.10.26.513817

**Authors:** Celeste Allaband, Amulya Lingaraju, Stephany Flores Ramos, Tanya Kumar, Haniyeh Javaheri, Maria D. Tiu, Ana Carolina Dantas Machado, Roland A. Richter, Emmanuel Elijah, Gabriel G. Haddad, Vanessa A. Leone, Pieter C. Dorrestein, Rob Knight, Amir Zarrinpar

**Affiliations:** Division of Biomedical Sciences, University of California, San Diego; La Jolla, CA, USA; Division of Gastroenterology, University of California, San Diego; La Jolla, CA, USA; Department of Pediatrics, University of California, San Diego; La Jolla, CA, USA; Medical Scientist Training Program, University of California San Diego; La Jolla, CA, USA; Department of Neurosciences, University of California, San Diego; La Jolla, CA, USA; Rady Children’s Hospital; San Diego, CA, USA; Department of Animal and Dairy Sciences, University of Wisconsin-Madison; Madison, WI, USA; Skaggs School of Pharmacy and Pharmaceutical Sciences, University of California, San Diego; La Jolla, CA, USA; Center for Microbiome Innovation, University of California, San Diego; La Jolla, CA, USA; Center for Computational Mass Spectrometry, University of California, San Diego; La Jolla, CA, USA; Department of Computer Science and Engineering, University of California, San Diego; La Jolla, CA, USA; VA Health Sciences San Diego; La Jolla, CA, USA; Institute of Diabetes and Metabolic Health, University of California, San Diego; La Jolla, CA, USA

## Abstract

Although many aspects of microbiome studies have been standardized to improve experimental replicability, none account for how the daily diurnal fluctuations in the gut lumen cause dynamic changes in 16S amplicon sequencing. Here we show that sample collection time affects the conclusions drawn from microbiome studies and are larger than the effect size of a daily experimental intervention or dietary changes. The timing of divergence of the microbiome composition between experimental and control groups are unique to each experiment. Sample collection times as short as only four hours apart lead to vastly different conclusions. Lack of consistency in the time of sample collection may explain poor cross-study replicability in microbiome research. Without looking at other data, the impact on other fields is unknown but potentially significant.

**One-Sentence Summary:** If we are not controlling for host circadian rhythm time in microbiome studies when performing experiments, it is like trying to measure sea level rise while not knowing that tides or waves exist.

## Main Text

The lack of replicability of microbiome studies has been a barrier to understanding how host-microbe interactions contribute to physiological homeostasis and pathophysiological processes, including heart disease and cancer^1^. As the field moves from descriptive and associative research to mechanistic and interventional studies, the ability to rapidly and reproducibly characterize the microbiome is critical to the development of novel microbiome-mediated therapeutics and diagnostic biomarkers^2^. In early studies, many confounding variables involving model systems, sample collection protocols, and pipeline processing were not routinely accounted for in study design, often resulting in irreproducible, noisy data^1,3^. The investigation of these irreproducible and noisy data led to the discovery of important confounds that influence the results, such as the maternal effect^4^, cage effect^5^, facility differences^6^, as well as laboratory and sample handling protocols^7^. However, despite the introduction of standardization of experimental protocols and analysis pipelines, unexplained variability and lack of replicability still plagues microbiome research.

One underexplored factor is that the microbiome is dynamic, and exhibits diurnal oscillations^8–10^. Disruption of microbiome diurnal dynamics^11–15^ are associated with metabolic syndrome spectrum diseases (e.g. insulin resistance, increased adiposity)^8^. A recent study found that an intestinal specific knockout of one of the circadian genes, *Bmal1*, in a mouse model was able to protect against diet-induced obesity^16^. The gut microbiome is intimately linked to host peripheral circadian rhythms and is known to influence physiology broadly, including behavior and thermoregulation^17^. Microbiome-depleted mice (i.e. antibiotic-induced depletion or germ-free mice) have dampened epithelial and hepatic circadian rhythms^11,18,19^. Analysis of the microbiome from human stool samples collected from a multitude of time points^20,21^, as well as 24-hour salivary collections^22–24^, suggest that the human microbiome also has diurnal fluctuations. In addition, loss of diurnal dynamics of the gut microbiome was recognized as a risk factor for developing type 2 diabetes in a longitudinal study of a large patient cohort^25^. Many labs that study the microbiome anecdotally report collecting their specimens for each experiment at a specific, single time point. However, it is not clear whether the collection time is chosen rationally based on experimental design, convenience to the experimenters, or if this window of time is consistent between experimental replicates both within and outside of the laboratory. We hypothesize that circadian variation is significant enough to affect microbiome results. By using existing diurnal microbiome studies, we can determine whether sampling at different times leads to different conclusions.

## RESULTS

### Time of sample collection is critical to microbiome study conclusions

To determine whether the time of sample collection was included in experimental methods of microbiome studies, we reviewed over 550 articles published in 2019 from major journals where new 16S or metagenomic datasets were generated. Only 0.32% reported a specific time of sample collection (**Fig S1A-C**). A recent study of biological sciences articles confirmed a low percentage of time-of-day information reporting in a broader field^26^. Since microbiome studies do not commonly report time of sample collection in their methods, we investigated the effects of microbiome sample collection on the potential interpretation of a study using the datasets from our meta-analysis. A targeted literature review followed by extensive correspondence, led to the acquisition of five previously published datasets in a form suitable for re-analysis (**Fig S1D**)^11,13,27–29^. In addition, we included a recently published dataset from the same mice in one of our circadian studies^13,30^. We also included analysis from an unpublished study that is unique in that it includes two circadian collections over the course of a single experiment. To quantify the effect of sample collection time on the microbial population, we used between-condition weighted UniFrac β-diversity distances (BCD) to show how similar microbiomes are to each other at any given time point. We chose weighted UniFrac because it takes into account both phylogeny and abundance of the organisms present. For circadian studies, standard notation of time of day is Zeitgeber time (ZT), where dawn/lights on is ZT0 (ex. our vivarium lights-on is 6AM, but this varies by facility). Thus, increasing or decreasing BCD allows us to assess microbiome compositional fluxes between experimental conditions over time.

First, we wanted to investigate whether sampling time affects the conclusion of a study with a discrete daily intervention. We started by reanalyzing a previous dataset ^27^ that used apolipoprotein E knock-out mice (*Apoe*^-/-^) mice under intermittent hypoxia and hypercapnia (IHC) conditions to mimic obstructive sleep apnea (**Fig 1A**). In the study, BCD fluctuated greatly, nearly doubling within a 24hr period (**Fig 1B**), suggesting that compositional assessments from different times would yield radically different results. Within-condition distances fluctuated much less during the same period (**Fig S2F, S2G**).

**Figure 1:**
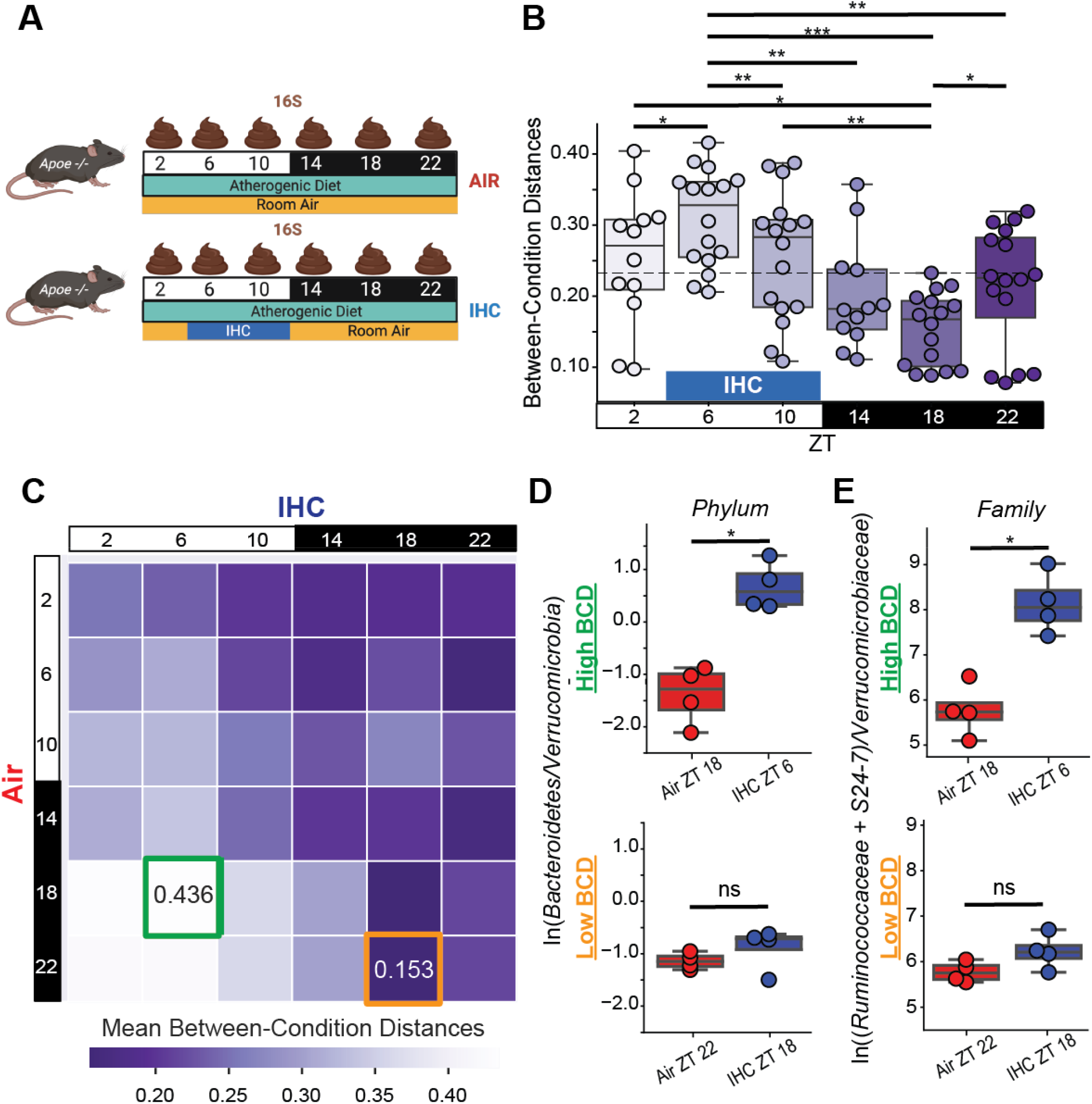
Microbiome Analysis of *Apoe*^-/-^ Mice Exposed to IHC Show Vastly Different Outcomes Depending on Time Point of Sample Collection. **A)** Experimental design. IHC= intermittent-hypoxia-with-hypercapnia (obstructive sleep apnea-like conditions). **B)** Between-condition distances (BCD), a subset of weighted UniFrac β-diversity distances. Significance is determined using paired Wilcoxon rank-sum test. The BCD values in this experiment were oscillating in a diurnal fashion (MetaCycle, JTK method, p<0.001). **C)** BCD heatmap by time point. Replicates were collapsed by taking the mean. Highest highlighted in green, lowest highlighted in orange. At the peak and trough time points identified in C, **(D)** the logarithmic ratios of differentially abundant key phyla of interest and **(E)** the logarithmic ratios of differentially abundant key families of interest. Notation: ns = not significant, * = p < 0.05; ** = p < 0.01; *** = p < 0.001

BCD increased during IHC exposure, with maximal divergence of the two groups at ZT-6 (Fig 1B, S2A, S2B). Maximal convergence (similarity) occurred at ZT-18, a half day after the maximal divergence when both groups were experimentally similar. Subsequently, despite the lack of the IHC intervention to separate the groups at that time, distances increased during ZT-22 which suggests a potential microbiome response to host anticipatory stress. In addition, the BCD values conformed to a diurnal pattern (MetaCycle, JTK method, p<0.001). Next, we used the distance matrix to create a heatmap of the average BCD between IHC and control mice for each time point combination to determine all potential outcomes of the study (max = 0.351; min = 0.082; range[max-min] = 0.269; mean = 0.232)(**Fig 1C**). The highest BCD (greatest divergence) between the two groups was Air ZT-18 and IHC ZT-6, which are 12 hrs apart. The lowest BCD (greatest convergence) between the two groups was Air ZT-22 and IHC ZT-18, both of which occur during the dark phase and are only 4 hrs apart. The highest BCD is 2.8 times the lowest across all timepoints, while the within-condition distances for Air (4.6X) and intermittent-hypoxia-hypercapnia (4.4X) dynamic ranges were greater (**Fig S2C-E**). The two groups had overall significantly different microbiome compositions (PERMANOVA, all Air vs all intermittent-hypoxia-hypercapnia, p=0.005), with ZT-6 driving differences (PERMANOVA, p=0.035). All other timepoints showed the two groups as being non-significantly different (PERMANOVA, p>0.05). Thus, the beta-diversity of the two conditions can differ 2.8-fold depending on the time of sample collection, potentially affecting the conclusions of the study.

To determine whether these different sampling times affect conclusions of the compositional analysis while accounting for bias caused by relative compositional bias and unknown microbial loads for each sample^31^, we examined log-ratios of biologically relevant phyla and families at the time points corresponding to the highest and lowest BCD (**Fig 1D**). The impact of sample collection time was most obvious at the phylum level, where the relative proportion of Bacteroidetes to Verrucomicrobia shifted strikingly towards Bacteroidetes in mice under IHC conditions at the highest BCD, but were indistinguishable at the lowest BCD. These differences existed at the sub-phylum level as well. For example, the log-ratio balance of three metabolically important families (Ruminococcaceae and S24-7, in relation to Verrucomicrobiaceae) shifted significantly during maximal BCD, but the balance was similar between experimental groups at the lowest BCD (**Fig 1E**). Thus, time of sample collection had a significant effect on microbiome composition and would have affected the conclusions made if only a single time point was performed.

### Diet and Feeding Pattern Influence Sample Collection Time Results

Since diet and feeding patterns induce large and reproducible effects on the gastrointestinal environment and resulting microniches ^32^, we hypothesized that it would be less influenced by diurnal microbiome dynamics.

We pursued this hypothesis by analyzing the results from one of our previously published studies ^13^ that investigated the effect of diet and feeding patterns on murine host physiology and the diurnal dynamics of the cecal microbiome. In mice on the same diet but with different feeding schedules, the BCD should change in response to differences in the feeding schedules of the experimental groups. In this experiment (**Fig 2A**), wild-type male C57Bl/6J mice were provided with either a normal chow diet (NCD) or a high-fat diet (HFD). Their access to food was either *ad libitum* (control; unrestricted access to food) or time-restricted (TRF). After 8 weeks, mice demonstrated a metabolic phenotype difference between HFD-*ad libitum* and HFD-TRF mice ^13^. Subgroups of mice were euthanized and the cecal contents were collected every 4 hours for 24hrs to examine dynamic changes in microbiome composition.

**Figure 2:**
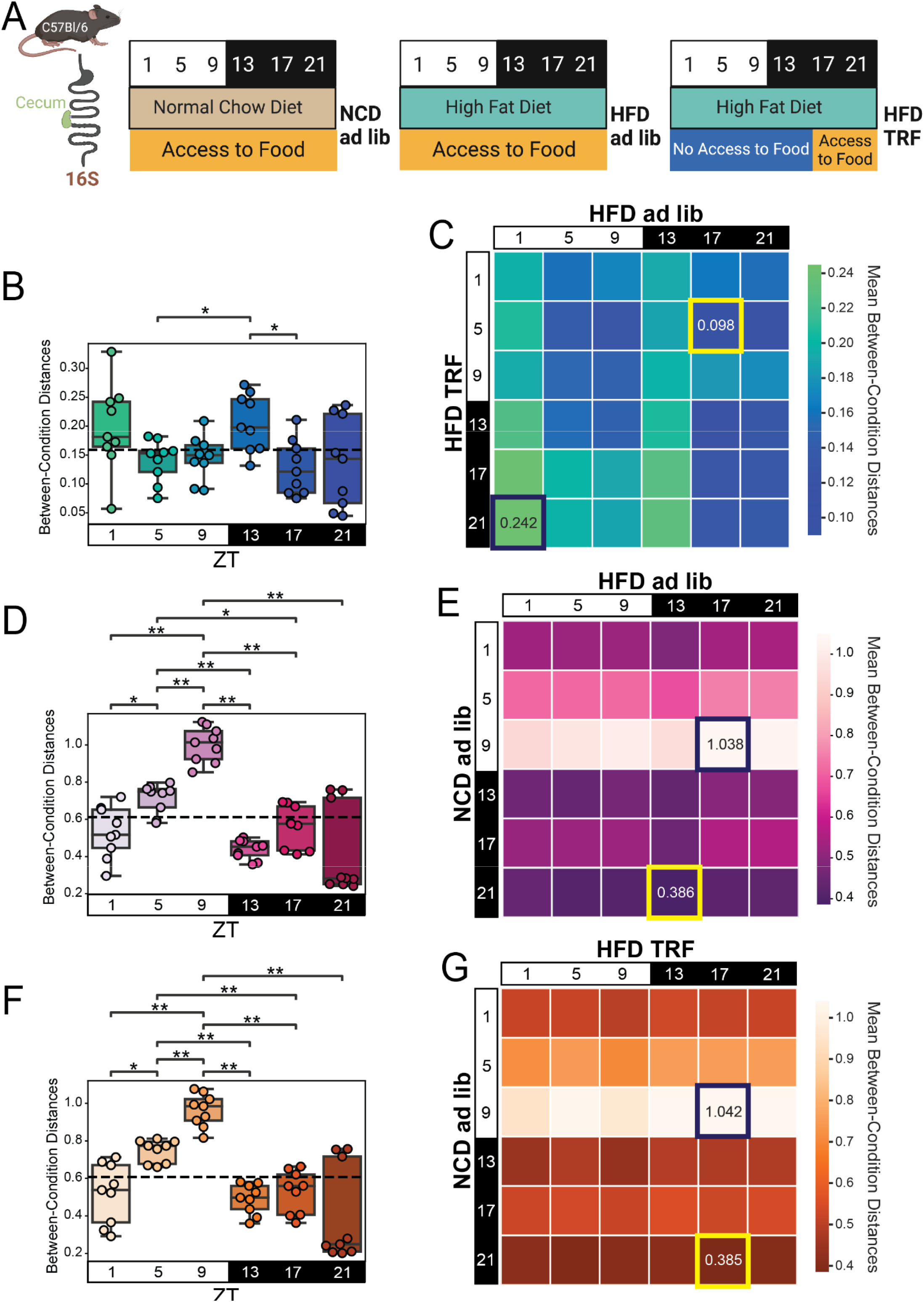
Diet and Feeding Pattern Influence Sample Collection Time Results in the Cecum. **A)** Experimental design. High Fat Diet (HFD). Time restricted feeding (TRF) - mice were restricted to eating only between ZT13-ZT21. Time point ZT13 was collected before the switch to eating, and thus mice were fasted at this time point. Time points were taken every 4 hrs for 24hrs (n=3 mice/condition/time point from separate cages; 6 total time points). **B)** BCD for cecal samples comparing HFD *ad libitum* vs HFD TRF. The dotted line is the average of all shown weighted UniFrac distances. Every point represents the calculated beta diversity distance between a control and experimental mouse. Within-group distances are shown in Figure S2. Significance was determined using a paired Mann-Whitney-Wilcoxon test two-sided with Bonferroni correction. **C)** Heatmap of cecal BCD between HFD *ad libitum* and HFD TRF mice by time point. Replicates were collapsed by taking the mean. Highest highlighted in indigo, lowest highlighted in yellow. **D)** BCD for cecal samples comparing NCD *ad libitum* vs HFD *ad libitum*. The dotted line is the average of all shown weighted UniFrac distances. Significance was determined using the Mann-Whitney-Wilcoxon test two-sided with Bonferroni correction. **E)** Heatmap of cecal BCD between NCD *ad libitum* controls and HFD TRF mice by time point. Replicates were collapsed by taking the mean. Highest highlighted in indigo, lowest highlighted in yellow. **F)** BCD for cecal samples comparing NCD *ad libitum* vs HFD TRF. The dotted line is the average of all shown weighted UniFrac distances. Significance was determined using the Mann-Whitney-Wilcoxon test two-sided with Bonferroni correction. **G)** Heatmap of cecal BCD between NCD *ad libitum* controls and HFD TRF mice by time point. Replicates were collapsed by taking the mean. Highest highlighted in indigo, lowest highlighted in yellow. Notation: * = p<0.05; ** = p<0.01; *** = p<0.001.

Since HFD mice spread their caloric intake throughout the day and night, we would expect low BCD between HFD-*ad libitum* and HFD-TRF mice during ZT-17 and ZT-21, when both groups have access to food. We expect high BCD during ZT-1 to ZT-13, when one group has access to food and the other group is forced to fast. As expected, the HFD-*ad libitum* to HFD-TRF BCD was the highest at ZT-13 when the two groups would be the most divergent (**Fig 2B**). We also saw that the HFD-*ad libitum* to HFD-TRF BCD was significantly lower at ZT-17. However, BCD was significantly lower at ZT-5 than ZT-13, and indistinguishable from ZT-17, suggesting that the intestinal environment is not solely influenced by the presence of a consumed diet in the lumen. Furthermore, the heatmap comparing all the combinations of different collection times shows a nearly 2.5-fold difference in peak and nadir BCD (max value = 0.242; min value= 0.098; range[max-min] = 0.144; mean = 0.159). There is a trend of the highest values being in the lower left corner (**Fig 2C**), which indicates that light phase of HFD-TRF and dark phase HFD*-ad libitum* have the greatest divergence. Thus, while the feeding schedule does impact microbiome composition, there are also composition shifts not directly attributable to the experimental design.

Next, we analyzed mice on different diets but with *ad libitum* access to food. Since diet macronutrient profile is a large driver of microbiome differences between cohorts, we wanted to determine if oscillatory dynamics of the gut microbiome could influence conclusions from microbiome compositional analysis. We hypothesized that the greatest differences between the two groups would be when they are eating different diets during the dark phase. Thus, we would expect the highest BCD to occur during ZT-13 to ZT-21 when one group is eating NCD and the other HFD. However, despite having radically different diets, the BCD from all of the dark phase time points are relatively low indicating similarities between the microbiomes (p>0.05)(**Fig 2D**). The biggest compositional shifts occurred at the transition from the light phase to the dark phase. The time point of greatest divergence is at ZT-9 when NCD mice are largely fasting, while HFD mice are likely eating at low to moderate levels. The heatmap shows a 2.7-fold difference in the peak to nadir BCD and also confirms that NCD-*ad libitum* ZT-9 as being different from all other HFD time points (max value = 1.038; min value= 0.386; range[max-min] =0.652; mean = 0.611)(**Fig 2E**). This same pattern is seen in a separate published dataset comparing NCD-*ad libitum* and an *ad libitum* milk-fat diet, that also yielded higher BCD during the light phase (mean = 0.416) and lower BCD during the dark phase (mean = 0.321) (**Fig S3**) ^11^. This indicates that the luminal environment differences caused by diet consumption alone do not drive differences between experimental groups and that dynamic oscillations of the luminal environment affect the interpretation of dietary changes, even with a powerful determinant of microbiome composition such as the macronutrient profile of the diet Next, we looked at a combination of both diet and feeding pattern differences, using NCD-*ad libitum* to HFD-TRF BCD. Since diet has such a huge effect on the microbiome, we hypothesized that the greatest differences between NCD-*ad libitum* and HFD-TRF would be when they are both eating different diets during the dark phase since both groups would be fasting during the light phase. Thus, we would expect the highest BCD to occur during ZT-17 or ZT-21. Opposite to our hypothesis, we found that the highest BCD values were during the light phase, especially ZT-9 (**Fig 2F**). Despite the two groups eating diets with vastly different macronutrient profiles we still saw a significant decrease in BCD values when we would have expected them to diverge. Thus, neither feeding/fasting rhythms nor diet alone drive these temporal fluctuations. In addition, the diurnal pattern of NCD-*ad libitum* to HFD-TRF BCD fluctuations most closely resembled the comparison between two different diets fed *ad libitum* (**Fig 2D**). The heatmap confirms a similar pattern of NCD-*ad libitum* to HFD-TRF value distribution across timepoints (**Fig 2G**) as NCD-*ad libitum* to HFD-*ad libitum* BCD (**Fig 2E**), with a 2.7-fold difference in peak to nadir BCD (max value = 1.042; min value= 0.385; range[max-min] =0.657; mean = 0.608). Since the mean BCD across timepoints is smaller than the maximum BCD at any timepoint minus the minimum BCD at any timepoint, the effect of sample collection time is thus larger than the effect size of a daily experimental intervention or dietary changes.

Thus, while the feeding pattern and diet do appear to significantly influence microbiome composition, their effects are not predictable on a timepoint-by-timepoint basis. Moreover, if an experimental variable effect as large and reproducible as that imposed by diet is affected by sample collection time, then experimental variables with smaller effects - such as medications, metabolites, and genotype - are likely to be even more variable with respect to time.

### Gastrointestinal Region Influence Sample Collection Time Results

Though the microbiome of the large and small intestine are quite different ^33^, the diurnal dynamics of the latter has only recently been characterized ^11,28,30^. We hypothesized that the dynamic response to changes in diet are not the same between gastrointestinal regions. We pursued this hypothesis by analyzing the results from a previously published study that investigated the diurnal dynamics between different GI regions ^11^. Leone, et al. compared a normal chow diet (NCD) to a high milk-fat diet (MFD) and examined the differences in the microbiome communities of both the cecum and ileum during a 24hr period (**Fig 3A**). The cecum and ileum had significantly different NCD-ad *libitum* to MFD-*ad libitum* BCD at ZT-6, in the middle of the light phase (**Fig 3B**). Thus, while microbial dynamics was generally similar between the two dietary conditions, there is at least one time point where time of sample collection would have made a difference when comparing dietary responses in the two organs. Heatmaps comparing BCD at different collection times for the ileal samples in this experiment show opposite trends in the timepoints of highest and lowest similarity compared to cecal samples (**Fig 3C, 3D**). While they had opposite trends in the timepoints that had the peak and trough values, the magnitude of change between these values was relatively similar with a 3.5-fold dynamic range in the cecum and 3.8-fold dynamic range in the ileum (**Fig 3C, 3D**). Thus, ileal and cecal diurnal dynamics are not always identical and can, at times, be significantly different.

**Figure 3:**
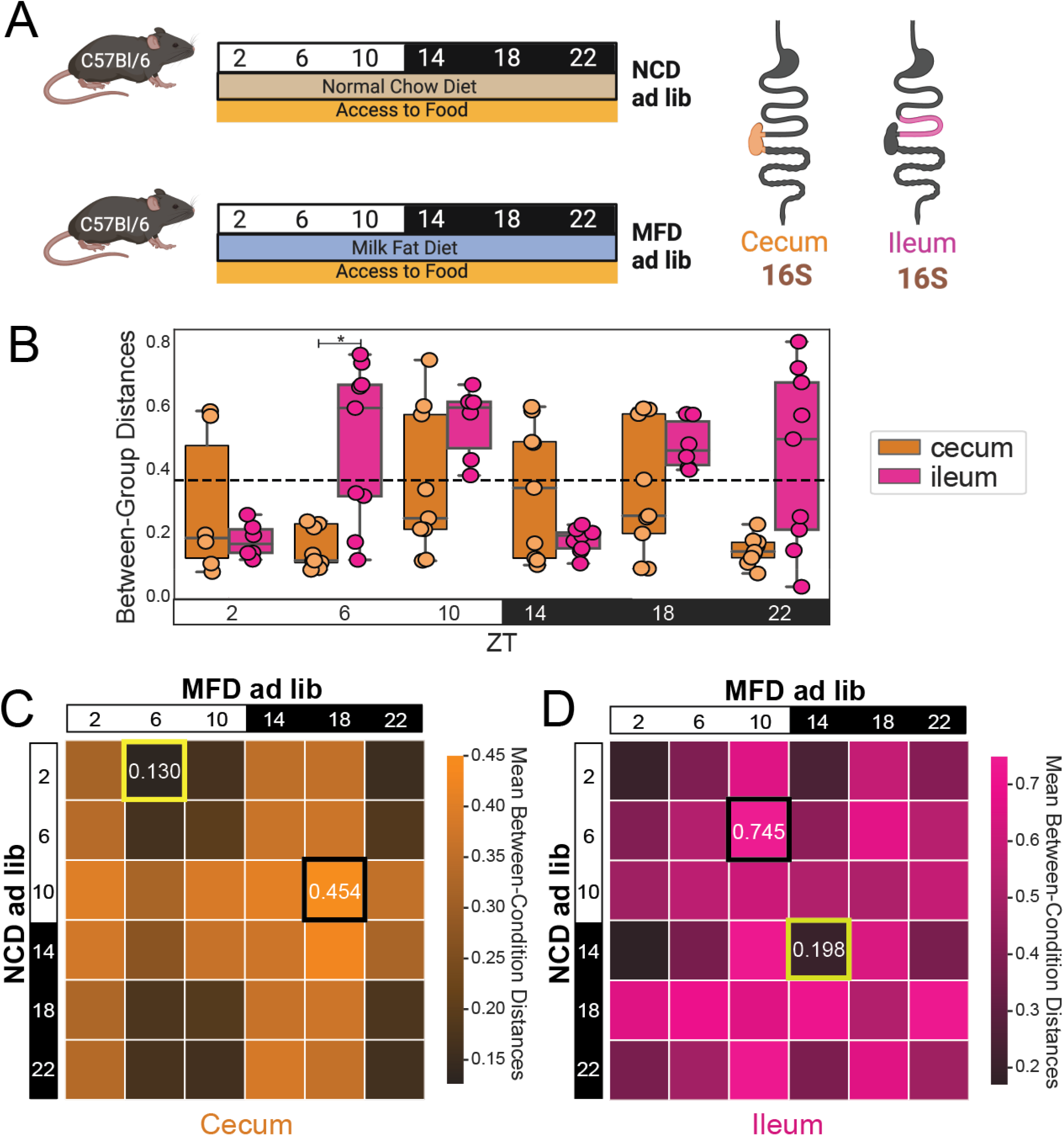
Gastrointestinal Region Influence Sample Collection Time Results. **A)** Experimental design. Mice were fed *ad libitum* with either normal chow diet (NCD) or high milk fat diet (MFD). After 5 weeks, cecal and ileal samples were collected every 4 hours for 24 hours (N=3 mice/condition/timepoint). **B)** BCD for both ileal and cecal samples comparing NCD vs MFD. The dotted line is the average of all shown weighted UniFrac distances. Ileal vs Cecal pairwise significance was determined using Mann-Whitney-Wilcoxon test two-sided with Bonferroni correction. **C)** Heatmap of BCD from cecal samples collected from NCD controls and HFD *ad libitum* mice by time point. Replicates were collapsed by taking the mean. Highest highlighted in black, lowest highlighted in yellow. **D)** Heatmap of BCD from ileal samples collected from NCD controls and HFD TRF mice by time point. Replicates were collapsed by taking the mean. Highest highlighted in black, lowest highlighted in yellow. Notation: * = p<0.05; ** = p<0.01; *** = p<0.001.

Moreover, the Zarrinpar, et al. study (used for the analysis presented in Fig 2) also had 16S rRNA amplicon sequencing from the ileum of the same mice that was recently published^30^. Similar to the Leone, et al. study, these results generally revealed different daily patterns in the ileum (**Fig S4**) than in the cecum (**Fig 2**). The dynamic range of values present in the heatmaps (highest BCD/lowest BCD) is approximately 3.0 in the ileum which is 15% higher than that in the cecum (**Fig 2C, E, G vs Fig S4C, E, G**). Thus these reproducible results show that the ileum responds differently over the course of the day than the cecum to the same conditions.

Finally, in a separate study, Wu, et al. investigated the effects of light exposure (i.e. 12h light:12h dark [LD] vs. 24hr dark [DD]) on the jejunal and ileal microbiome of Balb/c mice. The jejunal BCD was fairly consistent across all time points (**Fig S5**), suggesting that either this intervention (i.e. light exposure), or the proximal gut which has a more sparse microbiome, do not have the same dynamic shifts as the distal gut. Thus, though sampling time affects the outcomes studies on the ileal microbiome, it does not seem to affect the outcomes of studies in the jejunal samples. Furthermore, specific micro-niche sites (luminal and mucosal) within a single gastrointestinal region can have unique temporal patterns that are not expected based on experimental design alone (**Fig S6**). Together, these studies demonstrate different niches within the same mice have different microbiome dynamics.

### Longitudinal data is also susceptible to the influence of time

Samples from the IHC experiment (**Fig 1**) were collected a week after the experiment started with the intent to characterize the microbiome induced by the environmental exposure, prior to the dysmetabolic phenotype affecting the gut microbiome. However, in the TRF study (**Fig 2**), samples were collected after the phenotype was present. Since many microbiome experiments do not report the rationale for the timing of their sample collection, we questioned whether the length of experimental exposure time affects BCD. We performed a new study to examine where BCD changes over the course of a long study. In this study (**Fig 4A**), the *Ldlr* knock-out (*Ldlr*^*-/-*^) mice received either *ad libitum* (control group) or TRF (experimental group) access to the atherogenic diet. After 1 week (“early”; pre-phenotype development) and 20 weeks (“late”; post-phenotype development), we collected stool every 4 hours for 24hrs to examine dynamic changes in time point composition over the course of a long term experiment.

**Figure 4:**
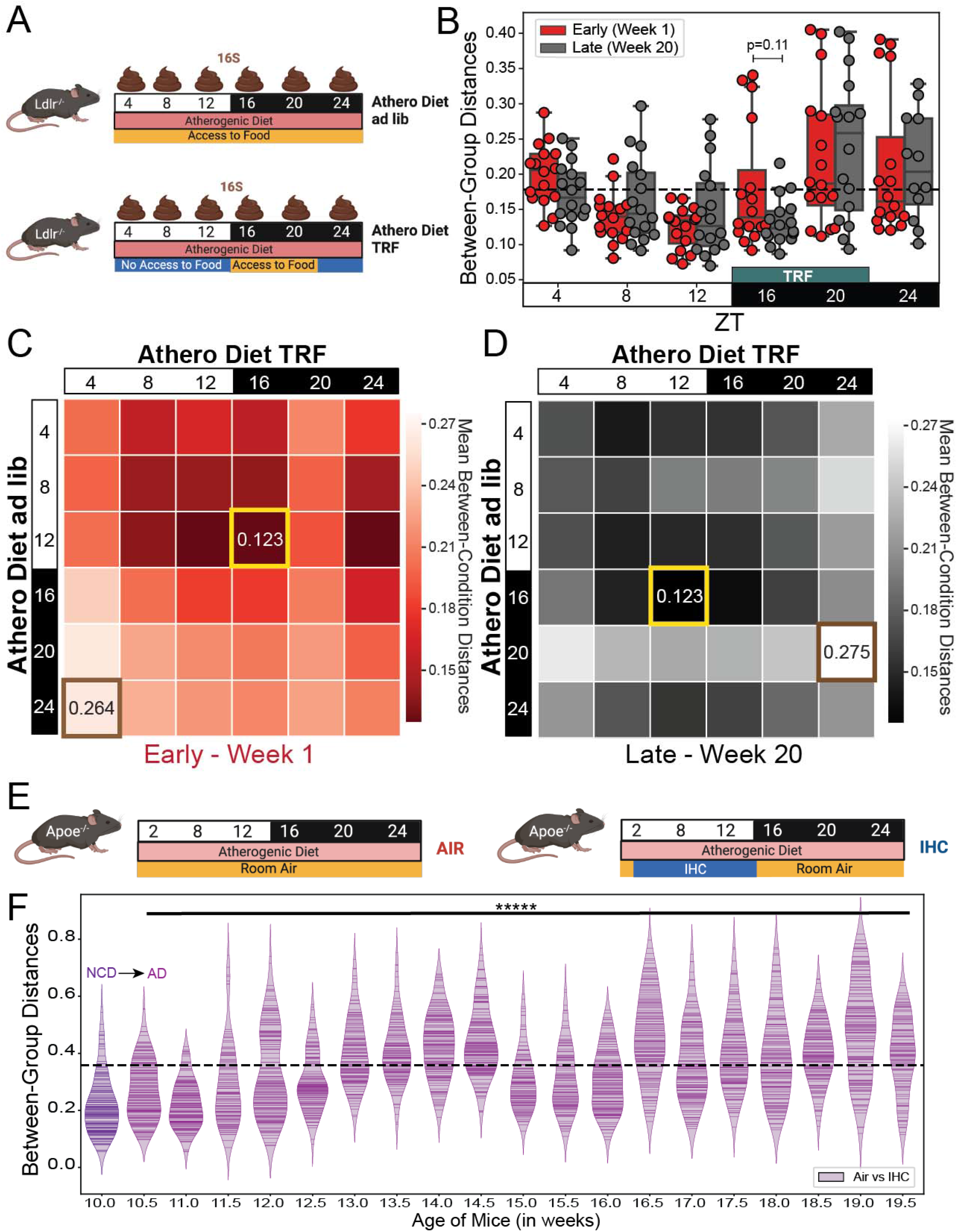
Longitudinal changes in BCD over the course of a study. **A)** Experimental design and sample collection for TRF study. Mice were fed atherogenic diet (AD) either *ad libitum* or TRF. Samples were collected every 4 hours for 24 hours (N=6 mice/condition/timepoint) after 1 week (early; pre-phenotype) and after 20 weeks (late; post-phenotype). **B)** BCD for *ad libitum* vs TRF conditions at the early (Week 1) and late (Week 20) timepoints. Dotted line is the average of all of the weighted UniFrac distances. Significance is determined using paired Wilcoxon rank-sum test. **C)** BCD heatmap for early samples, and **D)** BCD heatmap for late samples. Replicates were collapsed by taking the mean. Highest value is highlighted in tan and the lowest value is highlighted in yellow. **E)** Experimental design and sample collection for longitudinal IHC study. During the 10 weeks of exposure to either normal room air or IHC conditions, samples were collected between ZT-3 and ZT-5 every 3-4 days for the duration of the study (n=12 mice/condition).. **F)** BCD over the course of the IHC longitudinal study. Dotted line is the mean of all data shown. The only comparison shown is between Age 10.5 weeks and 19.5 weeks; significance was determined using paired Wilcoxon rank-sum tests. Notation: * = p<0.05; ** = p<0.01; *** = p<0.001, ***** = p<0.00001.

As shown in the previous studies, the time of sample collection during the day affects *ad libitum* to TRF BCD distances. During both the early and late phase of the experiment, maximum divergence *ad libitum* to TRF occurred during the dark period (highest mean BCD = ZT-20; **Fig 4B**). The BCD patterns conformed to a circadian-like pattern (p < 0.05, MetaCycle, JTK method) during both the early and late collection, with nearly identical amplitude and minor shifts in period and phase (**Fig S7**). Furthermore, the *ad libitum* to TRF BCD was not significantly different between the early and late part of the study at any time point **(Fig 4B**), demonstrating consistency within the study over time. The peak-to-trough ratios were also nearly identical between the early (**Fig 4C**) and late collection (**Fig 4D**). In general, these results demonstrate that longitudinal measures of BCD in a non-continuous intervention within a single experiment are relatively consistent over time.

To investigate the effects of longitudinal exposure to a daily discrete external intervention, we re-analyzed previously published data from our lab. In a previously published cohort of mice in an experiment investigating changes in the microbiome in response to IHC conditions (similar to **Fig 1A**) over several weeks until phenotype development (**Fig 4E**) ^34^. In this cohort, samples were collected once per day, during ZT-3 to ZT-5 (i.e. the time of greatest divergence), twice weekly over 10 weeks. While the control of IHC BCD fluctuated significantly during the course of the experiment, there was a slow generally upward trend (**Fig 4F**). The groups did diverge with significantly increased BCD over time (week 10.5 compared to week 19.5, p = 2.56×10^−8^, paired Wilcoxon rank sum, test statistic 1126) as the phenotype developed. Linear regression analysis resulted in a significant positive coefficient (p-value=6.72E-56, equation: y=0.016x+0.119). By holding the time of collection constant, we observed a compositional shift that occurred over time as the phenotype developed.

To determine if BCD is also relevant in longitudinal human studies, we re-analyzed a study that investigated the effects of a four day longitudinal dietary change (i.e. plant to animal based diet) in adult subjects on the speed and extent of shifts in the gut microbiome ^35^. When BCD was similarly calculated using weighted UniFrac, the plant-to-animal diet BCD demonstrated that the two groups did diverge the most on day 4 on condition (**Fig S8**). Since humans defecate on average once a day, it is difficult to investigate diurnal dynamics as we have done in mice. Recently, there have been attempts to reconstruct human diurnal rhythms using several thousand human samples ^25^, which have also shown diurnal pattern disruption in a disease state. Moreover, circadian rhythms are important from the beginning^36^ until the end^37^ of a human’s lifespan. Thus, time of sample collection is likely relevant in human samples as well.

## DISCUSSION

Since 2014 there has been unequivocal and reproducible research from multiple labs demonstrating diurnal fluctuations in the composition of the gut microbiome ^8,9,11–15,38,39^. Yet neither sample collection time nor the rationale for the selection of this time is reported outside of studies that are focused on diurnal fluctuations of the microbiome. Here, we show that the conclusions of a microbiome research study are greatly dependent on the time of sample collection, and that experimental and control groups undergo a cycle of diverging and converging microbiome composition depending on the nature and timing of experimental interventions. We hypothesize that host environmental differences at least partially drive these changes in gastrointestinal luminal microniches and cause divergence between the two conditions (BCD increase), converging again (BCD decrease) as the stress response fades.

Moreover, our findings suggest a fluidity of composition that is sensitive to a variety of host factors including environmental exposures, diet, gut region, and luminal micro-niche. Our BCD analysis confirms that, in some experiments, peak and trough distances can be as short as four hours apart. That is, shifting the collection of one condition by four hours could yield dramatically and potentially opposite conclusions on the similarity of the microbiome from experimental and control groups. This time scale may still be an overestimate; we did not collect stool samples at less than four-hour intervals. Thus, conflicting results from different laboratories may be due to differences in phase of the circadian cycle at the time of collection, timing relative to the experimental intervention, investigator chronotype (e.g. morning lark vs. night owl), or vivarium lighting setup. In studies with discrete daily interventions such as those described in this study, these differences can be quite pronounced. Based on our literature review, since the vast majority (>90%) of microbiome studies do not report when samples are collected, laboratories may unknowingly be collecting at suboptimal time points.

Furthermore, although it is likely a good assumption, due to convention and best utilization of researcher time, the methods section of published papers does not confirm that the control and experimental conditions are collected at the same time or within a specific window. In addition, while optimization of collection time points could be accomplished by sampling at the time of highest beta diversity for each group, caution should be taken not to artificially influence results. It would also be prudent to establish standard collection times for experiments in a field to ensure replication. To improve replicability, investigators should provide an explanation for the collection time of samples as it relates to their scientific hypothesis with the knowledge that anticipatory changes in the microbiome are quite pronounced.

While several of the studies used in this meta-analysis suffer from a low sample number, the fact that findings are replicated in laboratories from several different institutions, over several years, and with related study designs indicates this phenomenon is greatly understudied. Additional confounders can include changes in water content of stool due to time from defecation before sampling, which affects microbial density and richness as well as metabolism. Furthermore, to improve sensitivity to time, a study that attempts to deconvolute circadian rhythm and hours since sampling could be performed. While we have not had the opportunity to examine this phenomenon in metabolomics, viromics, and more, it is possible and even likely that circadian rhythms have impacts on these datasets as well. Examination of this phenomenon in humans is difficult because of infrequent defecation rates, but could potentially be recreated with large enough cohorts, such as KORA^40^, American Gut Project^41^, and FINRISK^42^. We propose that sample collection time be reported in ZT notation in future studies going forward. Otherwise, if we are not controlling for host circadian rhythm time, it is like trying to measure sea level rise while not knowing that tides or waves exist.

## Author Contributions

Conceptualization: CA, AZ

Methodology: CA, EE, PCD, RK, AZ

Investigation: CA, AL, SFR, TK, HJ, MDT, ACDM, RAR

Visualization: CA, SFR, TK, HJ, MDT, ACDM, RAR

Funding acquisition: AZ

Project administration: AZ

Supervision: RK, AZ

Writing – original draft: CA, AL, SFR, TK, HJ, MDT, AZ

Writing – review & editing: CA, AL, SFR, TK, HJ, MDT, ACDM, RAR, EE, GGH, VAL, PCD, RK, AZ

## Data Availability

### Microbiome

<u>Figure 1</u> - (Allaband/Zarrinpar 2021) - EBI accession <u>ERP110592</u>.

<u>Figure 2</u> - (Zarrinpar/Panda 2014) - see supplemental excel file attached to source paper [PMID: 25470548].

<u>Figure 3</u> - (Leone/Chang 2015) - The figshare accession number for the 16S amplicon sequence data: http://dx.doi.org/10.6084/m9.figshare.882928

<u>Figure 4</u> - (longitudinal circadian TRF) EBI: <u>ERP123226</u>; (longitudinal IHC) - EBI accession <u>ERP110592</u>.

<u>Figure S6</u> - (Tuganbaev/Elinav 2021) - ENA <u>PRJEB38869</u>

<u>Figure S8</u> (David/Turnbaugh 2012) - <u>MG-RAST</u> project ID 6248.

### Python Notebooks

https://github.com/knightlab-analyses/dynamics.

## List of Supplementary Materials

Materials and Methods

Fig S1 – S8

## Supplementary Materials

### Materials and Methods

#### Literature Review

(**Fig S1A-C**) We used the advanced search option from the four main journal groups, including the American Society for Microbiology (ASM) (https://msystems.asm.org), Science (https://search.sciencemag.org), Nature (https://www.nature.com), and Cell Press (https://www.cell.com). Searching for the term “microbiome” in all search fields (abstract, title, main text) during the year 2019 (Jan 1, 2019, to Dec 31, 2019) resulted in 586 articlesfrom 9 journals; mSystems (ASM), Science Translational Medicine (Science), Science Signaling (Science), Science Advances (Science), Science Immunology (Science), Nature (Nature), Nature Microbiology (Nature), Nature Communications (Nature), Cell Host Microbe (Cell), Cell (Cell), Cell Reports (Cell), Cell Metabolism (Cell). Our collection sheet includes a total of 16 columns: journal group, journal, year, article title, DOI, PMID, first author, last author, Microbiome (yes/no), vivarium (yes/no), vivarium setting, sample host, sample type, collection time, time note, and collection time reason. Notation of collection time was recorded as follows: explicitly stated (“yes”; 8AM, ZT4, etc.), implicitly stated (“relative”; “before surgery”, “in the morning”, etc.), or unstated (“not provided”; “daily”, “once a week”, etc.).

#### Systematic Review

(**Fig S1D**) When searching for the keywords “circadian microbiome” AND “mice” in PubMed (https://pubmed.ncbi.nlm.nih.gov/) for articles published over an 8 year period (from 2014-2021), we found 79 articles that met our initial criteria. Only 66 of those were research articles, and of the remainder we found only 14 articles that contained 16S amplicon sequencing samples collected for more than 3 time points within a 24 or 48 hour period. Of these 14 studies, four had complete publicly available data on ENA/EBI. Of the remainder, four had incomplete datasets on ENA/EBI - ^12,15,29,43^ - and the rest were not publicly available. We then contacted the authors of all studies with missing or incomplete data and got the following responses: four were unable to locate the missing data ^2,12,14,15^, three could not provide data in a format suitable for re-analysis ^43–45^, and three did not respond to repeated inquiries ^46–48^. This resulted in the acquisition of five previously published datasets in a form suitable for re-analysis ^11,13,27–29^.

#### Microbiome

All of the data in this paper is a re-analysis of previously published 16S studies, except for the data shown in Fig 4A-D (manuscript in preparation). Please refer to the respective source papers for detailed methods, including sample handling and preliminary processing. Raw data was procured from the respective data repositories as stated in the source paper, typically the European Nucleotide Archive (ENA). This data was then run through a standard QIIME2 pipeline (version 2021.8) ^49^ as follows: samples demultiplexed, denoised via deblur ^50^ into the amplicon sequence variant (ASV) table, feature table underwent rarefaction (as stated in source paper, see individual methods sections), representative sequences underwent fragment insertion on Greengenes_13_8 via SATé-enabled phylogenetic placement ^51^ to create the phylogenetic tree, and weighted UniFrac distances ^52^ were calculated. The resulting weighted unifrac distance matrix was filtered for only between-condition distances (BCD) as relevant to each study. Thus, using BCD values will show how similar the microbiomes from the two conditions are to each other at any given time point. Since BCD values are a subset of the Weighted UniFrac distance matrix values, both conditions (control and experimental) are taken into account with each distance value shown. Changes in BCD will demonstrate convergence (decreasing distance, increased similarity) or divergence (increasing distance, increased dissimilarity) of the microbiome composition between two groups. Circadian time notation is used throughout the paper to denote when samples were collected: Zeitgeber Time (ZT) were lights on = ZT-0. Data was visualized using custom python scripts, which can be found at https://github.com/knightlab-analyses/dynamics.

**Figure 1, S2** - Briefly, two groups of ten-week-old male *Apoe*^-/-^ mice on C57BL/6J background (002052; The Jackson Laboratory, Bar Harbor, ME) were individually housed in a 12-hour light:12-hour dark (12:12 L:D) vivarium. All mice were given an atherosclerotic-promoting diet (1.25% cholesterol, 21% milk fat; 4.5 Kcal/g; TD.96121; Envigo-Teklad Madison, WI) starting at 10 weeks of age until the end of the study. Mice in the experimental group were exposed to intermittent hypoxia and hypercapnia (IHC) conditions that consisted of 4 min of synchronized O_2_ reduction from 21% to 8% and synchronized elevation of CO_2_ from 0.5% to 8%, followed by alternating periods of 4 min of normoxia and normocapnia with 1- to 2-min ramp intervals. IHC conditions were administered in a computer-controlled atmosphere chamber (OxyCycler, Reming Bioinstruments, Redfield, NY) for 10 hours per day during the lights on phase (ZT-2 to ZT-12) when mice are sleeping for 10 weeks. Mice in the control group were exposed to normal room air (21% O_2_ and 0.5% CO_2_) during that same time period. After 6 days, fecal samples were collected every 4 hours for 24hrs (n = 4/group). 16S amplicon sequencing was performed on the V4 region using standard protocols (http://www.earthmicrobiome.org/emp-standard-protocols/). Rarefaction was set at 12,000 reads to control for sequencing effort. Please see the source paper for additional details ^27,53^.

**Figure 2, S4** - In short, wild-type SPF C57Bl/6 group-housed male mice (3 mice per cage) were provided either normal chow diet (LabDiet 5001, 13.5% calories from fat, crude fiber 5.1%) or a high fat diet (61% fat, HFD) and were fed in either an *ad libitum* manner, with access to food at all times, or fed in time-restricted (TRF) manner. TRF mice were allowed unrestricted access to HFD from ZT-13 to ZT-21. Mice on an NCD *ad libitum* diet (controls) typically fast during the light phase and consume >80% of their diet during the dark phase ^54,55^. However, mice on a HFD *ad libitum* diet (diet-induced obesity) lose this diurnal feeding pattern and spread their caloric intake throughout both the dark and light phase^54,55^. TRF of HFD consolidates feeding to the nocturnal period by providing access to food in a narrow time window, from ZT-13 to ZT-21 in this experiment, and is known to prevent the dysmetabolic effects of HFD consumption ^13,54,56^. After 8 weeks under these dietary conditions, mice were euthanized every 4 hours for 24hrs and intestinal contents collected (n=3 mice/condition/time point from separate cages; 6 time points). At ZT-13, fasted mice were euthanized prior to feeding. 16S amplicon sequencing was performed on the V1-V3 region using the 454 platform for cecal data. 16S amplicon sequencing was performed on the V4 region using Illumina primers for ileal data. For both regions, rarefaction was set to 1,000 reads to control for sequencing effort. Please see source paper for additional details ^13,30^.

**Figure 3, S3** - The study was performed on 8 to 10 week old male C57BI/6J SPF mice that were maintained in a 12:12 L:D cycle vivarium. The mice were fed *ad libitum* with either a normal chow diet (NCD, Harlan Teklad 2018S, 18% calories from fat, 3.5% crude fiber) or a 37.5% saturated milk fat diet (MFD, Harlan Teklad TD.97222 customized diet). <u>Figure 3</u> - After 5 weeks of being on the NCD or MFD diet, the mice were sacrificed and the cecal and ileal contents harvested every 4 hours for 24 hours (n = 3 mice/treatment); organ contents were flash frozen and stored at −80ºC. <u>Figure S3</u> - After 5 weeks of being on the NCD or MFD diet, fecal pellets were collected every 4 hours for 24 hours (n=3 mice/treatment); the fecal samples were stored at −80ºC. 16S amplicon sequencing was performed on the V4-V5 region using standard protocols (https://earthmicrobiome.org/protocols-and-standards/) in a High-Throughput Genome Analysis Core (Institute for Genomics & Systems Biology) at Argonne National Laboratory. Rarefaction was set at 10,000 reads to control for sequencing effort. Please see the source paper for additional details ^11^.

**Figure S6** - This study was performed on 8 to 12-week old WT C57BL/6 mice that were maintained in a 12:12 L:D cycle vivarium. The mice were fed a normal chow diet (NCD, Harlan Teklad 2018S, 18% calories from fat, 3.5% crude fiber) *ad libitum* for 4 weeks prior to sample collection. The mice were sacrificed and the luminal and mucosal small intestinal samples were collected every 4 hours for 24 hours (except for ZT-8, n = 4-5 mice/time point). The samples were frozen and stored at −80ºC. 16S amplicon sequencing was performed on the V4 region of the genome. Rarefaction was set to 4,200 reads to control sequencing effort. Please see the source paper for additional details ^28^.

**Figure 4A-D, S7** - This study was performed on 10 week old *Ldlr*^-/-^ mice (Jackson Labs) which were fed a high fat, high cholesterol diet (Research Diets D12109i; Clinton/Cybulsky high-fat rodent diet, regular casein, 1.25% added cholesterol, 0.5% sodium cholate). During the experiment, mice were maintained in 12:12 L:D reverse light-cycled cabinets (Phenome Technologies). Experimental and control groups were both on an atherogenic diet (AD), but one group was fed *ad libitum* and the other TRF. In TRF, mice were only allowed to eat for 8 hours per day during the dark phase of the day between ZT-13 and ZT-21. Fecal samples were collected every 4 hours for 24 hours (n=6 mice/condition) after 1 week (early; pre-phenotype) and after 20 weeks (late; post-phenotype). 16S rRNA was performed on the V4 region using the Earth Microbiome standard protocol (https://earthmicrobiome.org/protocols-and-standards/). Rarefaction was set at 11,498 reads to control for sequencing effort.

**Figure 4E-F** - In brief, two groups of ten-week-old male *Apoe*^-/-^ mice on C57BL/6J background (002052; The Jackson Laboratory, Bar Harbor, ME) were kept in a 12:12 L:D vivarium fed a normal chow diet (Teklad Rodent Diet 8604, 14% calories from fat, 4% crude fiber) before they were switched to an atherosclerotic-promoting diet containing 1.25% cholesterol and 21% milkfat (4.5 Kcal/g; TD.96121; Envigo-Teklad Madison, WI) starting at 10 weeks of age until the end of the study. Mice in the experimental group were exposed to IHC conditions as described in Fig 1 and were administered in a computer-controlled atmosphere chamber (OxyCycler, Reming Bioinstruments, Redfield, NY) for 10 hours per day during the lights on phase (ZT2-ZT12) for 10 weeks. Mice in the control group were exposed to normal room air (21% O_2_ and 0.5% CO_2_) during that same time period. Fecal samples were collected twice a week for the duration of the study ^34^.

**Figure S5** - In brief, two groups of five-week-old male *Balb/c* mice were kept in either a 12:12 L:D or 0:24 L:D vivarium fed a normal chow diet (unspecified in methods) *ad libitum*. After two weeks on condition, mice were anesthetized and sacrificed every 4 hours for 24hrs (n= 4-5 mice per group per time point). Samples from intestinal lumen, mucous layer, epithelial layer, and cecal contents were collected. The phenol-chloroform method was used for DNA extraction. 16S rRNA amplicon sequencing was performed on the V4 region. Rarefaction was set to 1085 reads to control sequencing effort, as performed in the source paper. Please refer to the source paper for detailed study design and associated protocols ^29^.

**Figure S8** - A total of 12 human subjects underwent 5 days of dietary intervention, either plant or animal based (n = 10 humans/condition). Patients that underwent both dietary interventions did so with a 1 month wash-out period in between interventions (10/12 patients; 9/10 patients per intervention). Two patients only underwent a single intervention (2/12 patients; one plant, one animal; 1/10). Please refer to the source paper for detailed study design and associated protocols ^35^.

## Supplementary Figures

**Supplemental Figure 1:**
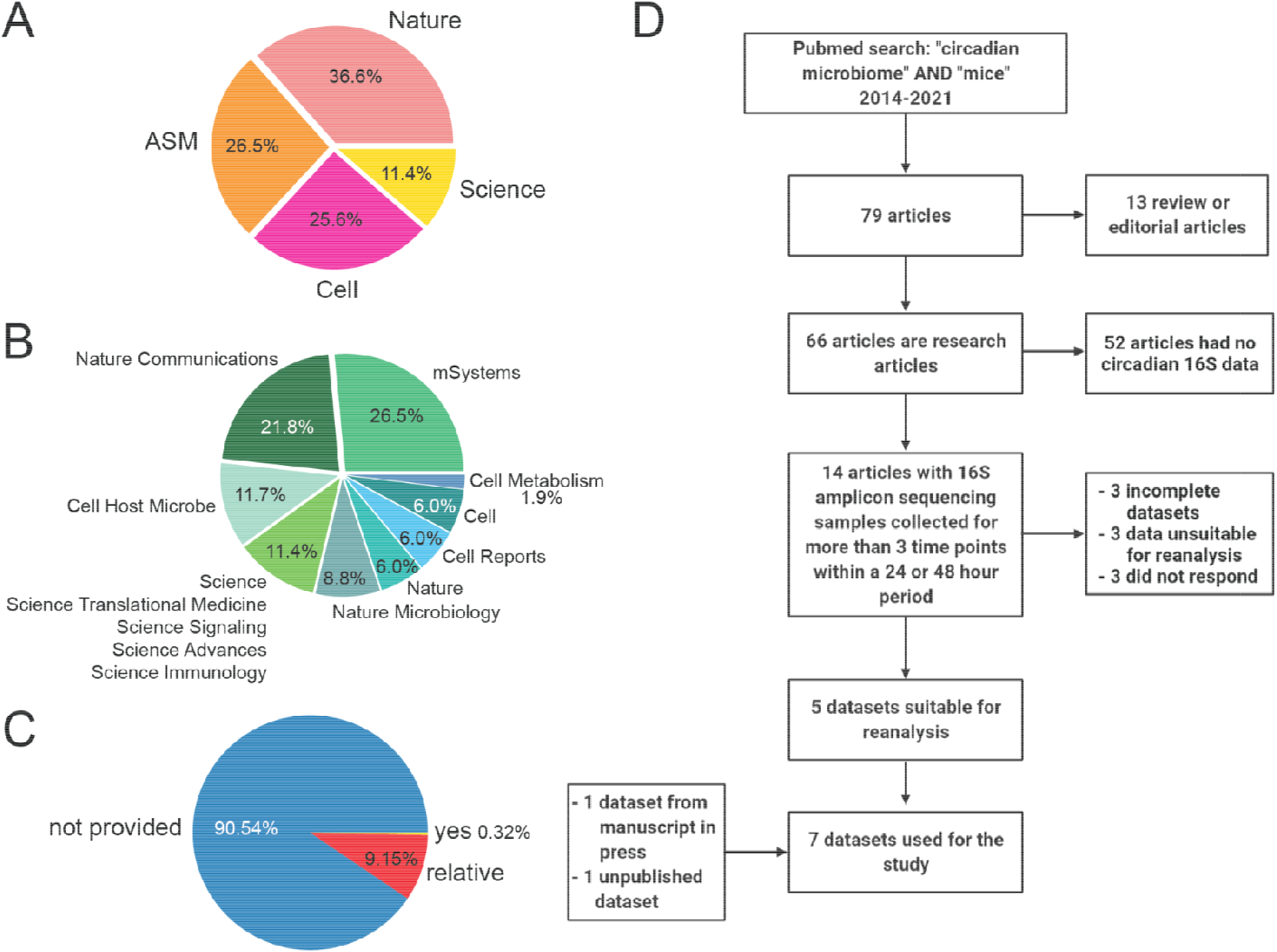
Circadian Microbiome Review. A) 2019 Literature Review Summary. Of the 586 articles containing microbiome (16S or metagenomic) data, found as described in the methods section, the percentage of microbiome articles from each of the publication groups. B) The percentage of microbiome articles belonging to each individual journal in 2019. Because the numerous individual journals from Science represented low percentages individually, they were grouped together. C) The percentage articles where collection time was explicitly stated (yes: 8 AM, ZT4, etc.), implicitly stated (relative: “before surgery”, “in the morning”, etc.), or unstated (not provided: “daily”, “once a week”, etc.). D) Meta-Analysis Inclusion Criteria Flow Chart. Literature review resulting in the five previously published datasets for meta-analysis ^11,13,27–29^.

**Supplemental Figure 2:**
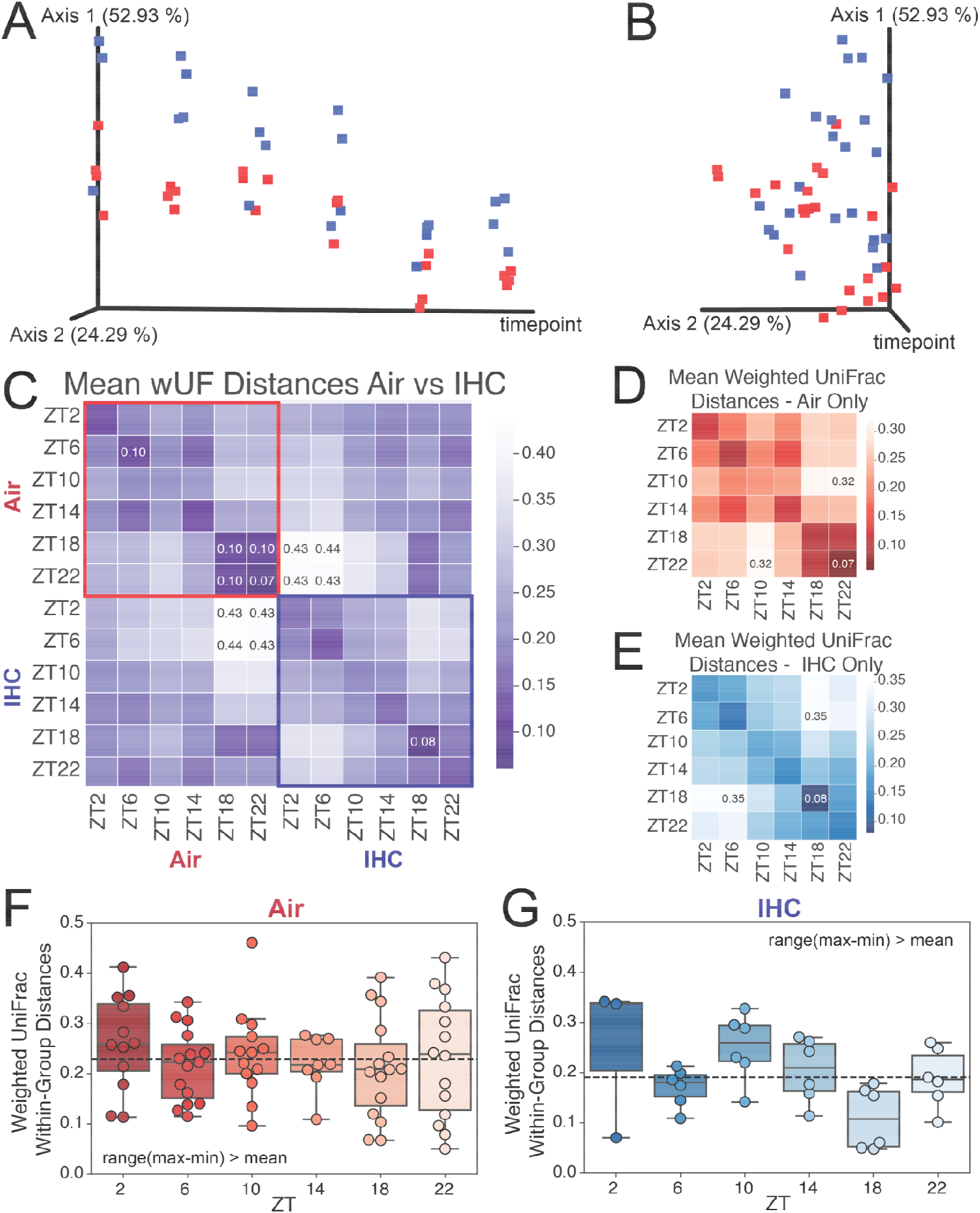
Diurnal IHC Weighted UniFrac PCoA and Within-Group Distances. A) Weighted UniFrac PCoA lateral view, with timepoints as one axis. B) Weighted UniFrac PCoA stacked view (same as A but different orientation) C) Full Weighted UniFrac distance matrix heatmap, averaged by timepoint. Red square indicates within-control group (Air) distances. Blue square indicates within-control group (IHC) distances. Top and bottom values labeled. D) Heatmap of mean weighted UniFrac distance values by timepoint, calculated using only control group (Air) samples. Top and bottom values labeled. E) Heatmap of mean weighted UniFrac distance values by timepoint, calculated using only experimental group (IHC) samples. Top and bottom values labeled. F) Boxplot/scatterplot of within-group weighted UniFrac distance values for the control group (Air). Zeros (ex. mouse1 ZT2 vs mouse1 ZT2 distance = 0) and duplicate values in the matrix were dropped. Dotted line indicates the mean of all values presented. No significant differences found. G) Boxplot/scatterplot of within-group weighted UniFrac distance values for the experimental group (IHC). Zeros (ex. mouse1 ZT2 vs mouse1 ZT2 distance = 0) and duplicate values in the matrix were dropped. Dotted line indicates the mean of all values presented. No significant differences found.

**Supplemental Figure 3:**
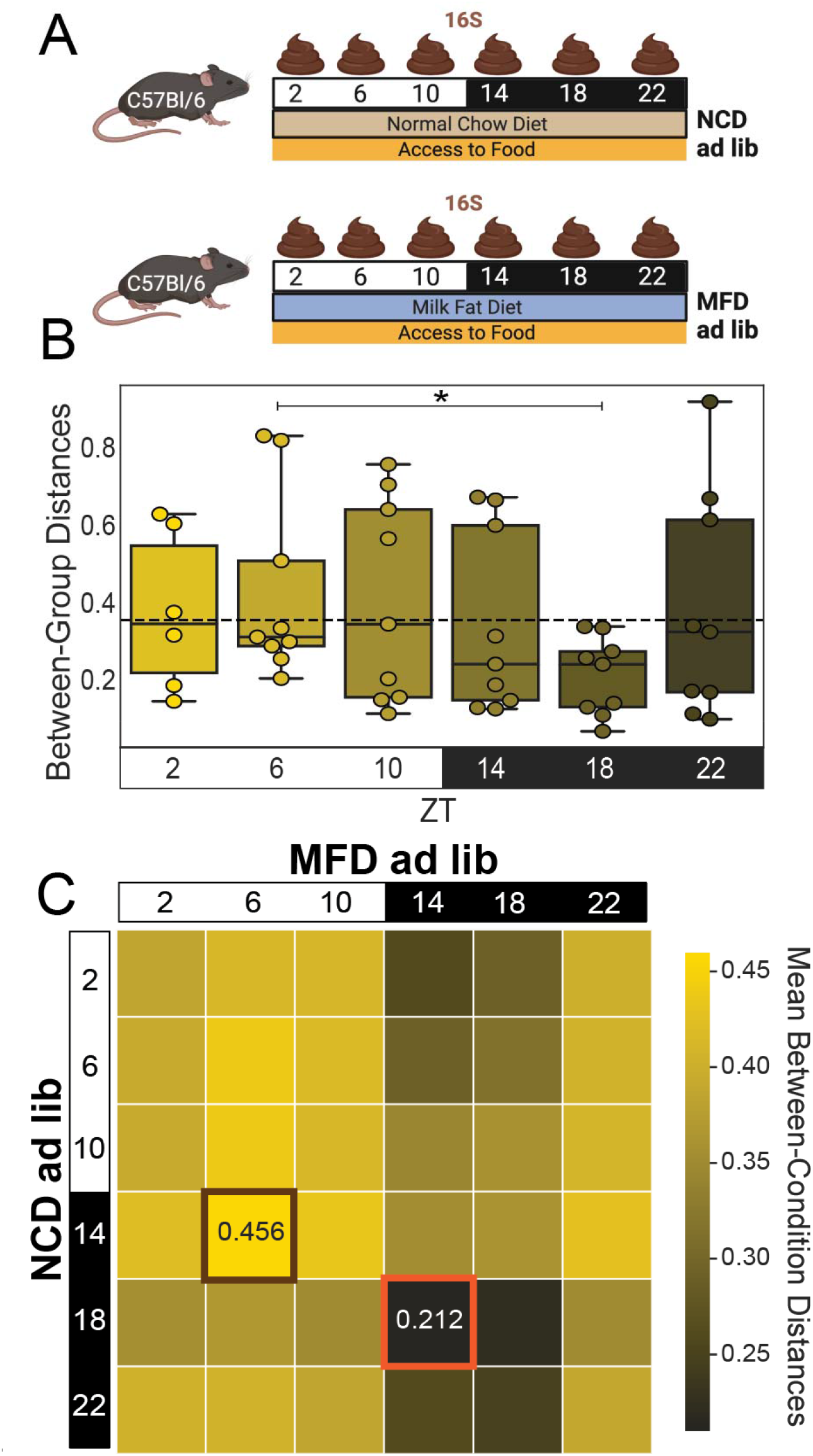
Temporal changes in BCD between NCD and MFD. A) Experiment design. C57Bl/6 mice were fed NCD (control) vs MFD ad libitum for 5 weeks before fecal samples were collected for analysis. Samples were collected every 4 hours for 24 hours (N=3 mice/condition). B) BCD for fecal samples comparing NCD vs MFD over 24hrs. The dotted line is the average of all shown weighted UniFrac distances. Significance was determined using the Mann-Whitney-Wilcoxon test two-sided with Bonferroni correction. C) Heatmap of mean BCD from fecal samples collected from NCD vs MFD mice by time point over 24hrs. Highest highlighted in brown, lowest highlighted in orange.

**Supplemental Figure 4:**
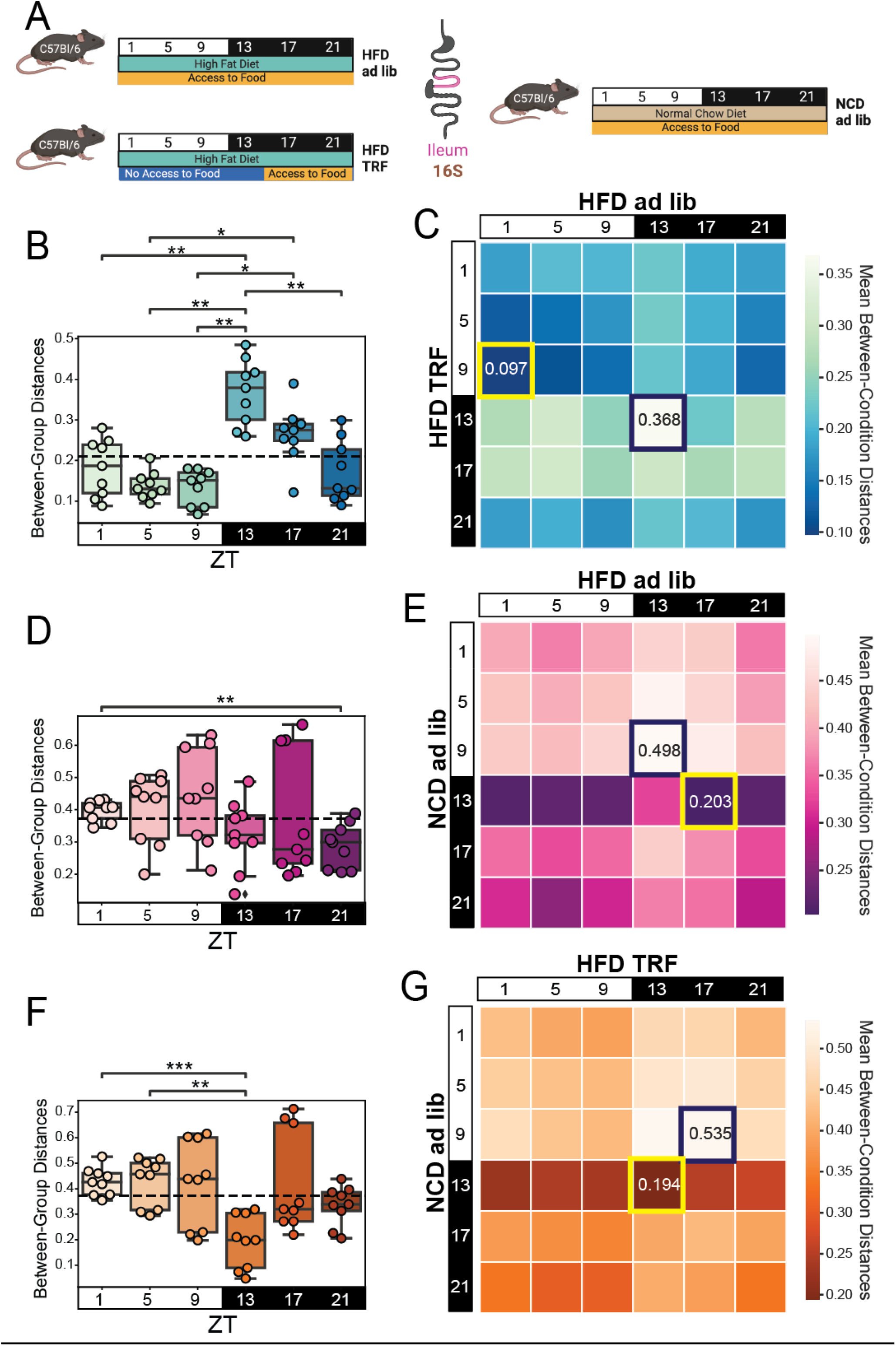
Diet and Feeding Pattern Influence Sample Collection Time Results in the Ileum. A) Experimental design. Mice used are the same as the ones in Fig 2 except this is unpublished ileal study. Mice were fed either ad libitum or TRF (ZT 13-21) access to HFD and compared to NCD ad libitum controls. After 8 weeks, ileal samples were collected every 4 hours for 24 hours (N=3 mice/condition). B) BCD for ileum samples comparing HFD ad libitum vs HFD TRF. Dotted line is the average of all shown weighted UniFrac distances. Significance was determined using a paired Mann-Whitney-Wilcoxon test two-sided with Bonferroni correction. C) Heatmap of mean BCD from ileum samples collected from NCD controls and HFD TRF mice by time point. Highest highlighted in indigo, lowest highlighted in yellow. D) BCD for ileal samples comparing NCD ad libitum vs HFD ad libitum. Dotted line is the average of all shown weighted UniFrac distances. Significance was determined using the Mann-Whitney-Wilcoxon test two-sided with Bonferroni correction. E) Heatmap of mean BCD from ileal samples collected from NCD controls and HFD TRF mice by time point. Highest highlighted in indigo, lowest highlighted in yellow. F) BCD for ileal samples comparing NCD ad libitum vs HFD TRF. Dotted line is the average of all shown weighted UniFrac distances. Significance was determined using the Mann-Whitney-Wilcoxon test two-sided with Bonferroni correction. G) Heatmap of mean BCD from ileal samples collected from NCD controls and HFD TRF mice by time point. Highest highlighted in indigo, lowest highlighted in yellow.

**Supplemental Figure 5:**
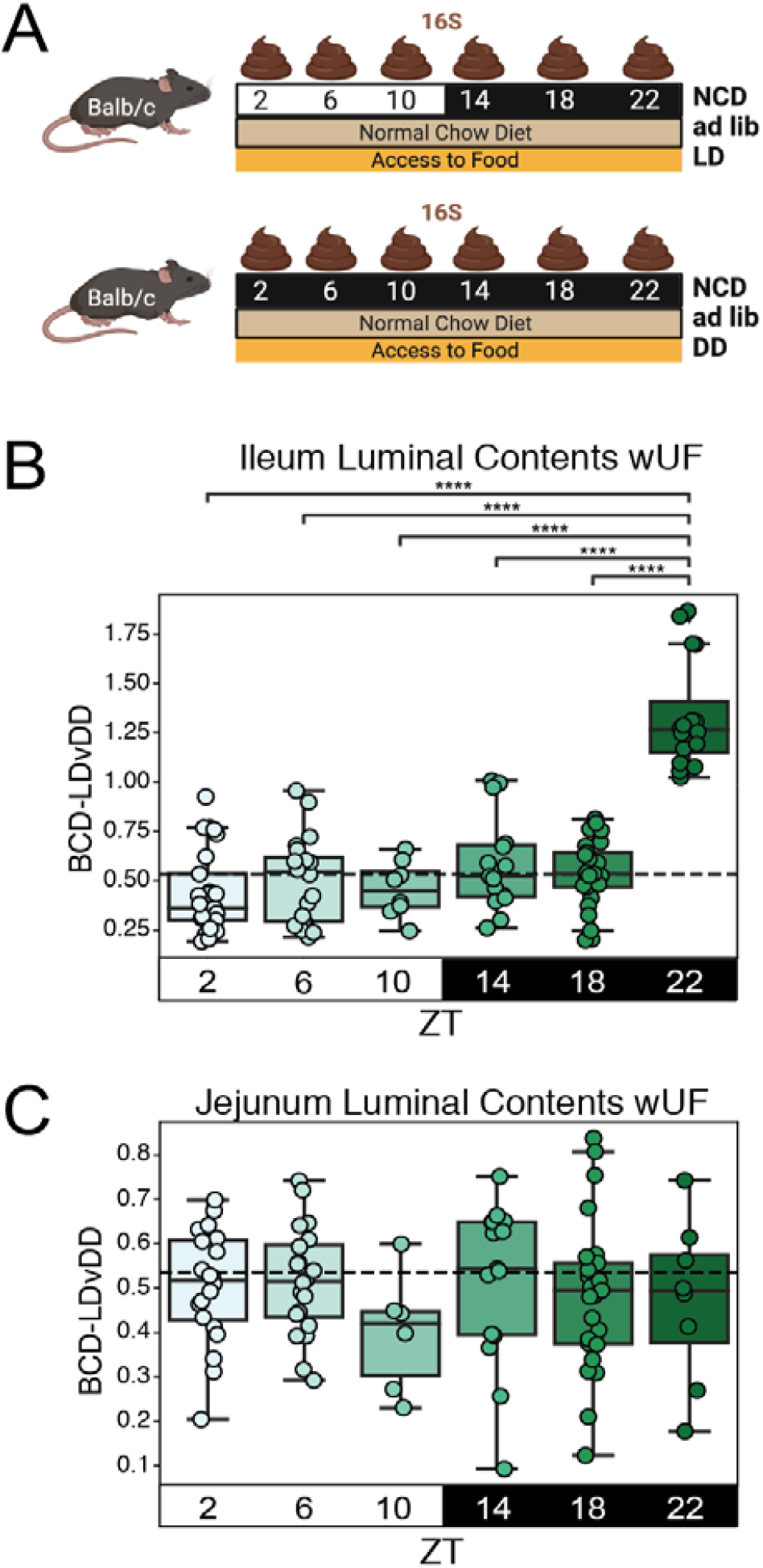
Irregular differences in diurnal rhythm patterns leads to generally minor shifts in BCD when comparing LD vs DD mice. A) Experimental design. Balb/c mice were fed NCD ad libitum under 0:24 L:D (24hr darkness, DD) experimental conditions and compared to 12:12 L:D (LD) control conditions. After 2 weeks, mice from each group were euthanized every 4 hours for 24 hours (N=4-5 mice/condition) and samples were collected from the proximal small intestine (“jejunum”) and distal small intestine (“ileum”) contents. B) BCD for luminal contents of proximal small intestine samples comparing LD to DD mice. Dotted line is the average of all shown weighted UniFrac distances. Significance was determined using a paired Mann-Whitney-Wilcoxon test two-sided with Bonferroni correction. C) BCD for luminal contents of distal small intestine samples comparing LD to DD mice. Dotted line is the average of all shown weighted UniFrac distances. Significance was determined using a paired Mann-Whitney-Wilcoxon test two-sided with Bonferroni correction.

**Supplemental Figure 6:**
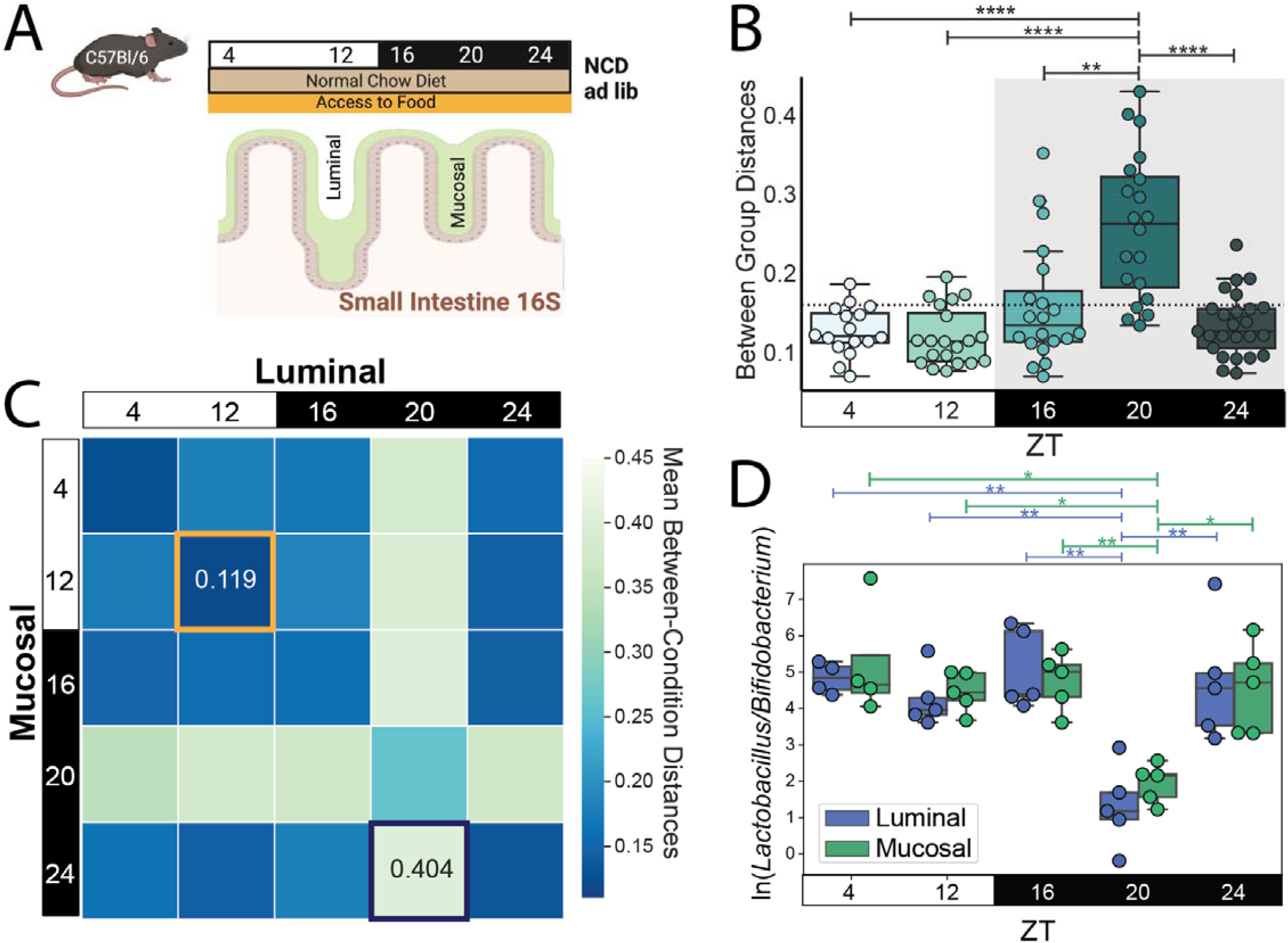
Localized changes in BCD between luminal and mucosal contents. A) Experimental design and sample collection for a local site study. Small intestinal samples were collected every 4 hours for 24 hours (N=4-5 mice/condition, skipping ZT8). Mice were fed *ad libitum* on the same diet (NCD) for 4 weeks before samples were taken. B) BCD for luminal vs mucosal conditions. The dotted line is the average of all shown weighted UniFrac distances. Significance is determined using the Mann-Whitney-Wilcoxon test two-sided with Bonferroni correction. C) Heatmap of mean BCD distances comparing luminal and mucosal by time point. Highest value highlighted in navy, lowest value highlighted in gold. D) Experimentally relevant log ratio, highlighting the changes seen at ZT20. Significance was determined using paired Wilcoxon rank-sum test. Notation: * = p<0.05; ** = p<0.01; *** = p<0.001, ***** = p<0.00001.

**Supplemental Figure 7:**
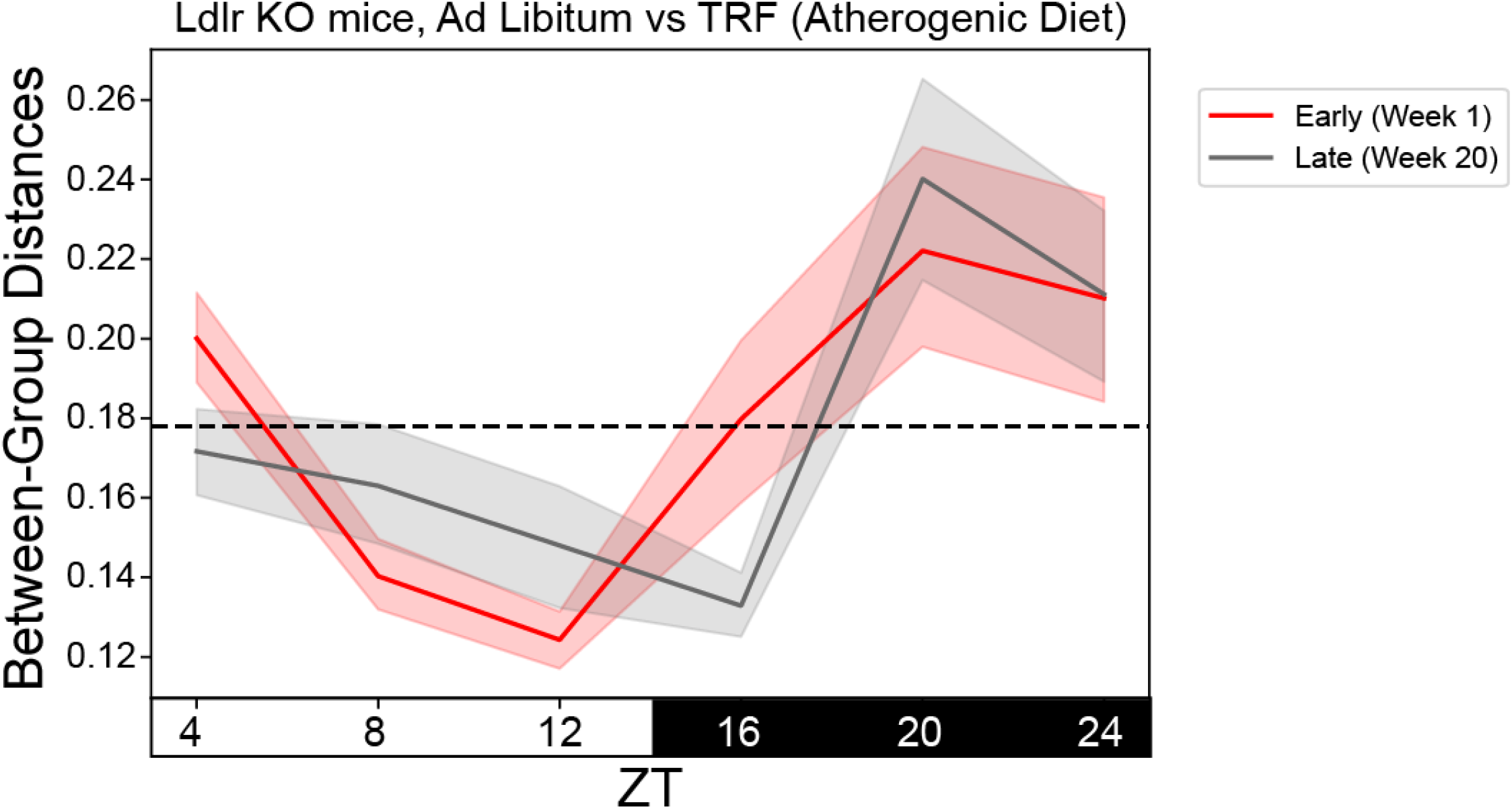
Line plot of the same values presented in Fig 3B. A)The shaded region represents the standard error of the mean. The dotted line is the average of all of the weighted UniFrac distances used to calculate this plot. Some of the shifts seen between early and late values may be more easily visualized in this format. <u>Early data</u>: MetaCycle, meta2d/LS method, p=0.0017, amplitude=0.215, period=22.3, adjphase=18.0; <u>Late Data:</u> MetaCycle, meta2d/LS method, p=0.0128, amplitude=0.198, period=25.7, adjphase=16.2.

**Supplemental Figure 8:**
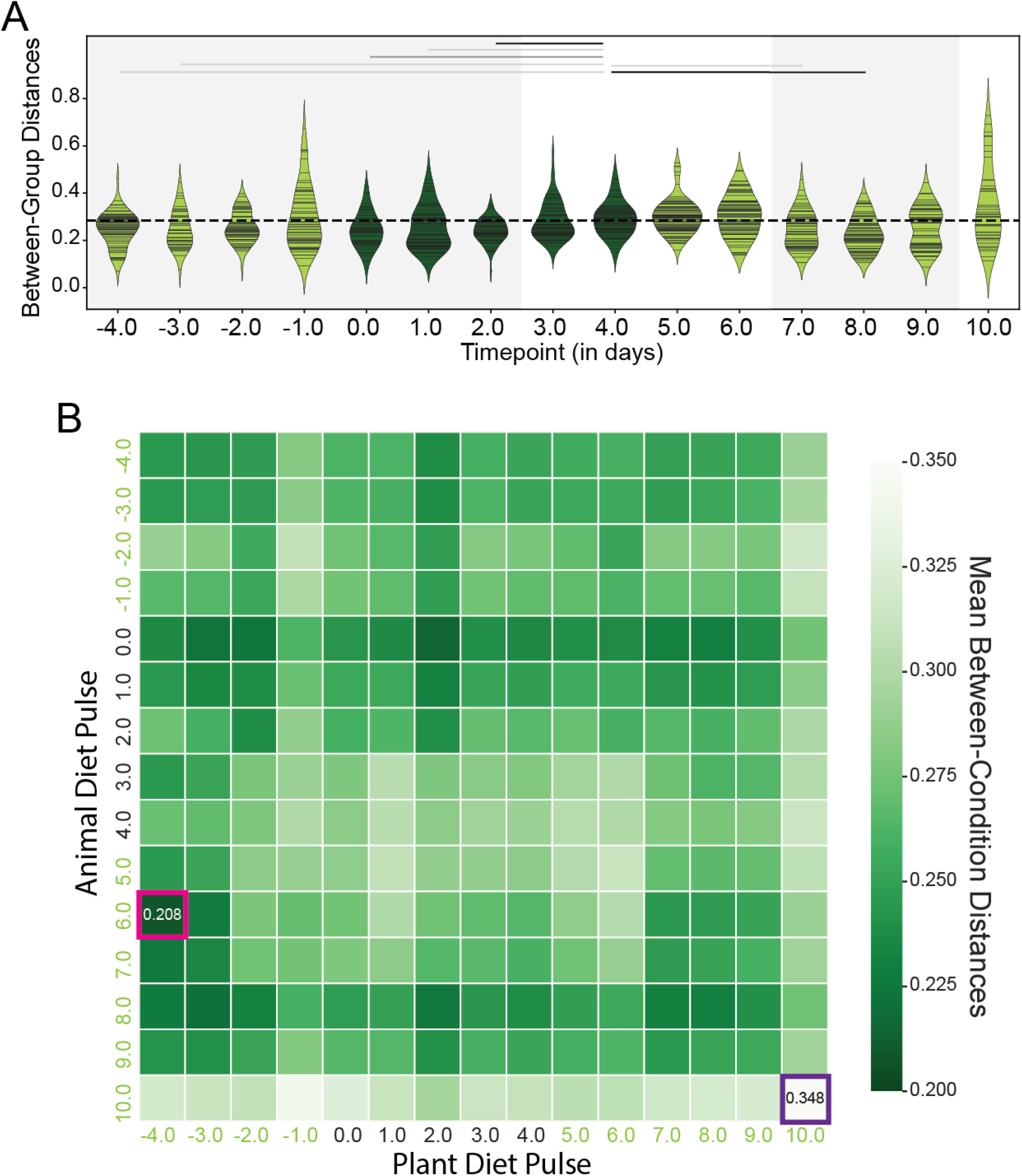
Human data also shows that a non-continuous intervention affects beta diversity distances over the course of a study. Experimental design: The patients underwent 5 days of dietary intervention, either plant or animal-based (N=10 humans/condition): 9/10 patients underwent both dietary interventions after a 1 month wash-out period, 1/10 patients only underwent a single intervention. See reference (36). A) Weighted UniFrac β-diversity violin plot using between-group distances for plant and animal dietary interventions. Each line on the violin plot is a sample value. The dotted line is the average of all of the weighted UniFrac distances from the time points farthest from the intervention (−4.0 and 10.0). The shaded area represents time points that are not significantly different from each other, except as noted. Significance was determined using the Mann-Whitney-Wilcoxon test two-sided with Bonferroni correction. Notation: light gray line = p<0.05; medium gray line = p<0.01; black line = p<0.0001. B) Mean weighted UniFrac β-diversity distance heatmap using values calculated between plant and animal dietary interventions by time point. Highest value highlighted in purple, lowest highlighted in pink.

